# Colchicine Therapy Provides No Functional or Anti-inflammatory Benefit in a Porcine Myocardial Infarction Model

**DOI:** 10.64898/2026.11.25.769313

**Authors:** Rebekka Grampp, Anja Stadtmüller, Florian Schnitter, Finja Becker, Thorsten Bischler, Panagiota Arampatzi, Nadine Gladow, Niklas Beyersdorf, Wolfgang R. Bauer, Anna Frey, Ulrich Hofmann

**Affiliations:** University Hospital Würzburg, Department of Internal Medicine I, Würzburg, Germany; University Hospital Würzburg, Comprehensive Heart Failure Center, Würzburg, Germany; University of Würzburg, Institute for Virology and Immunobiology, Würzburg, Germany; Department of Cardiovascular Imaging, Chair of Molecular and Cellular Imaging, Comprehensive Heart Failure Center, University Hospital Würzburg, Würzburg, Germany; Core Unit Systems Medicine, University of Würzburg, 97080 Würzburg, Germany

**Keywords:** Myocardial infarction, pig, colchicine, inflammation

## Abstract

**Aims:** We aimed to study whether deferred short-term colchicine treatment attenuates myocardial inflammation and preserves systolic function in a porcine myocardial infarction (MI) model.

**Methods and Results:** We induced MI in a total of 19 female landrace pigs by occlusion of the left anterior descending artery for 90 minutes. Animals were randomized to receive oral colchicine (0.05 mg/kg body weight) or placebo once daily from days 3–7. All 13 animals that received treatment survived until day seven. In both treatment groups, no left ventricular thrombi were detected. Cardiac function was assessed by echocardiography before MI and after 7 days. The mean change in ejection fraction (EF) was -25±19% in the colchicine and - 16±7 % in the placebo group (p= 0.36). Infarct size was 24±6% in the colchicine and 26±5 % in the placebo group (p= 0.13). On day 7, we determined infarct size by planimetry, myocardial leukocyte composition by flow cytometry and IL-1-beta (IL-1β) in infarct tissue by an ELISA assay. RNA-sequencing of heart tissue was performed for exploratory analyses of transcriptional changes imposed by colchicine treatment. Colchicine neither changed the leukocyte composition in the infarcted heart nor significantly reduced myocardial IL-1β protein levels or transcriptional patterns relating to proinflammatory activity in the heart.

**Conclusion:** The chosen colchicine treatment regimen was safe but did not improve EF or reduce infarct sizes. Inefficiency to attenuate inflammatory processes in the infarcted myocardium seems to be the most plausible cause for the observed lack of efficacy.

## 1. Introduction

Colchicine is a clinically long-established alkaloid derived from *Colchicum autumnale*. Its potent anti-inflammatory properties have been used for decades in the treatment of gout. Colchicine accumulates in white blood cells and decreases their motility, chemotaxis and NLRP3 inflammasome activity, thereby reducing pro-inflammatory cytokine release and the potential ensuing tissue damage(1). Several clinical trials have demonstrated its efficacy for reducing ischaemic events in patients with chronic coronary artery disease and in those following an acute MI(2, 3). Based on these trials, colchicine is approved for reducing ischaemic vascular events in patients with coronary heart disease.

Besides beneficial effects on atherosclerotic disease, experiments in rodents have indicated that colchicine mitigates cardiac ischaemia–reperfusion injury and preserves systolic function(4–7) when administered before or right after initiation of reperfusion. It is therefore tempting to speculate that early initiation of colchicine treatment after MI could help to limit both cardiac injury and recurrent vascular events. Encouraged by these promising preclinical results, the COVERT-MI trial tested the efficacy of colchicine in reducing infarct size and preserving ventricular function by initiating oral treatment before or immediately after reperfusion(8). The clinical trial revealed the unanticipated finding that colchicine treatment did not reduce infarct size but was associated with the formation of left ventricular thrombi. In the recent CLEAR-SYNERGY trial(9), initiation of colchicine treatment within 72 hours did not confer protection against ischaemic events although no unanticipated safety issues emerged. Moreover, data from a post hoc analysis of the COLCOT trial revealed, that there was a significant reduction in the incidence of the primary endpoint in patients in whom colchicine had been initiated within the first 3 days after MI compared to placebo without any apparent safety concerns(10). Therefore, while colchicine administration during the peri-reperfusion period may increase the risk of left ventricular thrombus formation, delayed initiation of therapy appears to be safe in patients with acute MI. Nevertheless, the optimal timing for treatment initiation remains uncertain. Specifically, it remains to be determined how early colchicine can be administered without compromising safety, while preserving its potential to limit infarct size, improve systolic function, and favourably modulate post-infarction healing. As neither the CLEAR-SYNERGY(9) nor the COLCOT trial(3) revealed safety concerns for patients treated with colchicine early after MI, initiation of treatment within the first few days after MI, sparing the early reperfusion phase, may represent the most appropriate therapeutic window. Nevertheless, the reasons underlying the unexpected absence of efficacy of colchicine in limiting myocardial injury or preserving systolic function, as observed in both the aforementioned porcine and clinical studies, remain to be elucidated.

Therefore, we aimed to characterise the effect of an oral colchicine treatment regimen on the resolution of early pro-inflammatory activity after MI, which is a key process for proper healing, alongside with its impact on infarct size and myocardial function. Using several complementary and orthogonal approaches to characterise post-infarction inflammation in a porcine MI model, we found no evidence of significant suppression of pro-inflammatory processes in response to colchicine treatment.

## 2. Methods

### 2.1 Animal experiments

Female German Landrace obtain from Gerd Heinrichs (Heinsberg, Germany) were maintained in the animal facility of the Comprehensive Heart Failure Center Würzburg at the University Hospital of Würzburg. All animal experiments were conducted in accordance with the German Animal Protection Law and had been reviewed and approved by the review board of the District Government of Lower Franconia and the University of Würzburg (approval reference number 55.2.2.-2532-2-1852).

A total of 20 pigs with a weight of 20-30 kg and an age of 8 ± 2 weeks were sedated by intramuscular injection of the neuroleptic azaperone (Stresnil® 40 mg/ml, Elanco GmbH, Germany or Sedanol® 40 mg/ml, Wirtschaftsgenossenschaft Deutscher Tierärzte eG) was administered at a dose of 8 mg/kg BW together with atropine (atropine sulphate 0.5 mg/ml, B. Braun Melsungen AG, Germany, dose 0.05 mg/kg BW). As soon as the effect of the sedation became noticeable after a few minutes, 15 mg/kg BW of ketamine (Ursotamine® 100 mg/ml, Serumwerk Bernburg AG, Germany) was administered i.m..

Then a venous catheter was placed in the lateral auricular vein (V.) on both sides. This was followed by spraying lidocain on the glottis region and performing endotracheal intubation and transfer to the operating room with spontaneous breathing.

Deepening and maintenance of anesthesia was achieved by continuous intravenous infusion of 5 mg/kg BW/h propofol (Narcofol 10 mg/ml, CP-Pharma Handelsgesellschaft mbH) and 0.012 - 0.024 mg/kg BW/h fentanyl (Fentadon 50 µg/ml Dechra Veterinary Products Deutschland GmbH).

In addition, a permanent intravenous infusion of 2 mg/kg BW/h amiodarone (Cordarex®, 150 mg/3 ml, Sanofi - Aventis Deutschland GmbH) in 5% glucose solution (B. Braun injection solution, B. Braun, Melsungen GmbH, Germany) for antiarrhythmic purposes and 10 ml/kg BW/h Sterofundin® (Sterofundin® ISO 1/1 E ISO, B. Braun Melsungen GmbH, Germany) were administered to maintain euvolemia. After cessation of spontaneous breathing, the animal was connected to mechanical ventilator (Primus®, Drägerwerk AG & Co. KGaA, Germany) (pressure control, percentage of inhaled oxygen in total volume 60%) for controlled ventilation. During anaesthesia, the animal’s vital parameters were permanently monitored: continuous, non-invasive monitoring of peripheral blood pressure, pulse rate and oxygen saturation, body temperature, expiratory CO_2_ and ECG data (patient monitor M3046A M3, Philips GmbH, Germany). After inserting an arterial sheath into the femoral artery for MI induction, its arterial blood pressure was monitored invasively. Echocardiography and sequencing data were analyzed by investigators blinded for treatment allocation.

### 2.2 Induction of myocardial infarction

The anesthetized pig was placed in supine position. The femoral artery was punctured under steep cautery and sonographic visualization, and the 6 French sheath was inserted into the vessel using the Seldinger technique. 225 IU/kg BW of heparin (Heparin Sodium 25000-ratiopharm®, 25,000 IU/5 ml, Ratiopharm GmbH, Germany) were administered intravenously to prevent thrombus formation during cardiac catheterization. First, the guidewire (EMERALDTM Guidewire, 175cm, 0.035’’, PTFE coating, J-tip 3mm, fixed, Cordis GmbH, Germany) was inserted into the deflated catheter (Launcher Guiding Catheter 6 F .071” Relaxed Mod, HSIISH, Medtronic plc, Ireland) and both were pushed through the sheath under X-ray guidance (Exposcop 8000, Ziehm imaging GmbH, Germany). The guidewire was first advanced via the ascending aorta to the aortic arch and the catheter followed via this. Once the catheter was in the left atrium, the wire was removed and the catheter was positioned so that the coronary sinus aorta could be visualized using a contrast agent.

A coronary wire (Hi-Torque Balance Middleweight Universal III 0.014” 190 cm, Abbott GmbH, Germany) was pre-bent to fit and pushed over the catheter far distally into the paraconal interventricular ramus. This served as a guidewire for the deflated balloon catheter (Emerge monorail 3.50 mm x 15 mm, Boston Scientific Medizintechnik GmbH, Germany), which was positioned distal to the first lateral branch and filled with contrast medium to approx. 6 mbar using a hand pump (Encore® 26 Advantage Kit, Boston Scientific Corporation, USA). This occluded the vessel and its occlusion was controlled by administering a bolus of contrast medium. Occlusion was maintained for a period of 90 minutes and could be tracked using ST segment elevation. Any ventricular fibrillation that occurred could be immediately detected via ECG monitoring and counteracted by defibrillation (bisphasic, 200 J) and cardiopulmonary resuscitation. In cases of asystole, adrenaline was administered intravenously (1 mg every 3–5 minutes) combined with chest compressions to maintain perfusion. If the above measures did not result in the restoration of spontaneous sinus rhythm, the animal was euthanized after prolonged resuscitation (max. 10 minutes) by intravenous bolus administration of pentobarbital (45-90 mg/kg BW).

After successful 90-minute occlusion, coronary angiography was performed both before and after balloon deflation to confirm reperfusion of the vessel. The coronary wire, balloon, and catheter were then removed, and the puncture site in the groin was closed using a closure system (Exoseal®; Cordis, Miami Lakes, FL, USA). This was followed by intramuscular administration of meloxicam (0.4 mg/kg BW) for postoperative pain relief, followed by emergence from anesthesia and extubation. The recovery phase was closely monitored. Meloxicam was administered again the following day.

Of the 19 animals one animal had to be euthanized prior to catheterization due to supraventricular tachycardia. 18 animals underwent the complete MI procedure, 6 died during coronary occlusion due to malignant arrhythmias. The surviving animals were sacrificed on day 7 post MI.

### 2.3 Echocardiography

Animals were examined both on day 0 before the induction of myocardial infarction (MI) and on day 7 post-MI, before euthanasia. For the ultrasound examination by an GE Vivid E9 echocardiography machine, the animal was connected to the ventilator while breathing spontaneously. Anaesthesia was maintained by intravenous administration of bolus doses of propofol. The examination was performed in both right and left lateral positions, starting with the right lateral position and using a transducer oriented from subxiphoid to cranial.

Subsequently, the animal was moved to the left lateral position, and both the long and short axes were imaged from a parasternal view. In the short axis, cross-sections of the heart at different levels can be displayed. Two and four chamber views were recorded but not included in quantitative analysis as proper image acquisition was not possible in all animals. The quantitative analysis based on long-axis M-modes was performed in a post-processing manner by a blinded researcher not involved in the animal experiments.

### 2.4 Organ processing

Finally, after administering a bolus of heparin (300 IU/kg BW) the animals were euthanized by injecting pentobarbital (Release® 500 mg/ml, WDT Garbsen, Germany) 45 (>30kg BW) – 90 (<30kg BW) mg/kg BW, into the ear vein.

After the heart was removed and washed with ice-cold 0.9 % NaCl to remove any remaining blood, it was transferred to the laboratory in cooled isotonic 0.9 % NaCl solution. Blood was also taken in two different tubes: one with heparin to prevent clotting and one without additive (serum tube) to obtain serum.

Both atria and right ventricles were separated along the septum, leaving only the left ventricle. The left ventricle was dissected into thin slices in order to subsequently examine specific areas in detail, such as infarct core (IC), border zone (BZ) and healthy myocardium (remote myocardium, R). Two 300 mg tissue samples were taken from each of these areas and transferred to 1.5 mL Eppendorf reaction tubes (Safe-Lock Biopur® Tubes 1.5 mL, Eppendorf, Germany) containing Hanks’ Balanced Salts solution (HBSS+/+, with Ca & Mg, w/o Phenol Red, sterile filtered, Capricorn Scientific, Germany). Using small surgical scissors, the heart material was minced within the reaction tubes and then transferred with cold HBSS +/+ into 50 ml Falcons (tube 50ml, 114 x 28 mm, DNA-, DNase-, RNase-free, Sarstedt AG & Co. KG, Germany), which after the addition of DNase I (60 units / ml, source: bovine pancreas, 10104159001, Sigma Aldrich/ Roche Diagnostics GmbH, Mannheim, Germany) and collagenase type 2 (600 units / ml, source: Clostridium hydrolyticum, LS004176, Worthington Biochemical Corporation, Lakewood, NJ, USA) are topped up with HBSS +/+, which was heated to 37°C in a water bath before, and placed in the thermal shaker (ThermoMixer®, Eppendorf, 4 x 50 ml,) for 30 minutes of digestion. After 30 minutes, digestion was stopped by adding cold HBSS/BSA (Hanks’ Balanced Salt Solution without calcium, without magnesium, with sodium hydrogen carbonate, RNBM0561, Merck Sigma-Aldrich, SIGMA-ALDRICH CHEMIE GmbH, Germany with specially added BSA, bovine serum albumin/albumin from bovine serum, heat shock protein, protease-free, pH 7, Sigma-Aldrich, A3294-50G) and the sample material was stained through cell strainers with a mesh size of 70 µm, followed by cell strainers with a mesh size of 40 µm. For a further purification of the sample, Red Blood Cell-lysis buffer (room temperature, RBC –lysis Buffer, 10x, BioLegend, San Diego, CA, USA) was added and after 8 minutes washed off with DPBS EDTA (Dulbecco’s Balanced Salt Solution without Ca2+/Mg2+, containing 2 mM ethylene diamine tetra-acetic acid) in room temperature. After the pellets have been resuspended in DPBS EDTA they were stained with 1:200 viability dye Zombie Aqua^TM^ (Zomobie Aqua^TM^ Fixable Viability Kit (DMSO), BioLegend, San Diego, CA, USA) and incubated for 15 minutes at a dark place with room temperature. Thereafter followed another washing step with flow cytometry buffer (DPBS−/−/EDTA with 2% fetal calf serum) and blocking of the cells with pig serum 10% (own production; in DPBS−/−/EDTA) for 15 minutes at 4 °C.

The cells were then stained with anti-CD45-FITC antibody (1:50, Clone K252.1E4; Isotype IgG1, Biorad, Hercules, CA, USA) and kept at 4°C for 30 minutes. After washing the excessive antibody off with FACS buffer (PBS/EDTA + 2% FCS) the pellets were resuspended in MACS (Magnetic activated cell sorting) buffer (PBS + EDTA + 0,5% BSA), mixed with MicroBeads (1:20, Miltenyi Biotec Anti-FITC, Bergisch Gladbach, Germany) and again stored for 15 minutes at 4°C, followed by a washing step with MACS buffer. The buffer ensures optimal conditions during the separation process and contributes to the efficiency and accuracy of cell separation, which is done by using LS columns (Miltenyi Biotec, Bergisch Gladbach, Germany) and due to the previously magnetically labeled cells also a strong magnet (QuadroMACS^TM^ Seperator, Miltenyi Biotec, Germany). The cell suspension obtained for each area of the heart was then divided for the different staining panels. The respective antibody mixtures have been prepared earlier and contain specific antibodies for myeloid and lymphoid staining. The cell suspension of each area was divided and stained with the corresponding antibody mix, whereby an unstained sample and additionally several FMO samples were examined during establishment. After staining with the antibody mixture, the cells were placed at 4°C for 30 minutes.

### 2.5 Flow cytometry

Antibodies used for staining leukocytes are listed in the Supplemental Table 1. Two panels were stained for characterizing leukocyte subsets:

Lymphoid: After the antibody mix (CD25, CD172a, CD8α, CD8β, CD3ε, CD21, CD4α) was added, the cells were incubated for 30 minutes and then washed. They were permeabilized by the addition of FixPerm (eBioscience Foxp3/Transcription Factor Staining Buffer Set - Thermo Fisher Scientifc, Carlsbad, CA, USA) for 45 minutes at room temperature. This step was necessary for the following intracellular FoxP3 staining. After 45 minutes at room temperature the cells were finally washed, resuspended with FACS buffer (including DPBS, EDTA, 2% FCS) and stored at 4°C until measurement.

Myeloid: The antibody mix (CD203a, CD163, CD172α, SWC8) was followed by staining with the secondary antibody (1:200 IgM Monoclonal Antibody (II/41) PE-Cyanine5; eBioscience™; Invitrogen) and incubation at 4°C for 30 minutes. Similar to the lymphoid panel the cells were also incubated with FixPerm (45 minutes, room temperature) but then stained with an intracellular antibody mix consisting of GPNMB (PE) and NLRP3 (AF647). After 45 minutes at room temperature, the cells were finally washed, resuspended with FACS buffer (DPBS + EDTA + 2% FCS) and acquired within the next two hours. Until then they were stored at 4°C.

### 2.6 Flow cytometric immunophenotyping

The stained cells were measured using the Invitrogen™ Attune™ NxT Flow Cytometer (Thermo Fisher Scientific, Waltham, MA, USA), and further analyses were performed using FlowJo v10.10.0 (FlowJo Single Cell Analysis Software, Ashland, OR, USA). Singlets were selected from all events by plotting FSC-A against FSC-H. Within the population of living cells (negative for Zombie Aqua), further subpopulations were identified from CD45-positive cells (e.g., leukocytes). For gating strategy see Suppl. Fig.I and II

### 2.7 Isolation of whole blood leukocytes

10 ml of fresh, heparinized blood of every animal in the trial from day 0 (MI) and day 7 (organ harvesting) were used for the isolation of whole blood leukocytes. A cryoring was thawed and filled with isopropanol up to the indicated line. Then, it was pre-cooled in a 4°C refrigerator. For hypotonic lysis, 40 mL of distilled water were added to 8 mL of heparinized whole blood and gently slewed. The reaction was stopped by adding 10x PBS and after centrifugation at 4°C, the supernatant was removed, the pellet resuspended Ery-Lysis buffer (at room temperature) and incubated for 8 minutes. Then, BSS/BSA (cold) was added in excess, centrifuged again and the remaining pellet (after removing the supernatant) resuspended in 10 mL of cold DPBS. The next step was to make a 1:100 dilution (990 µL PBS + 10 µL of the cell solution) of which 20 µL were pipetted onto a slide for the BioRad Cell Counter (BioRad TC10^TM^ Automated Cell Counter, Life Science Group). To freeze the cells at a concentration of 10 million cells per mL, the cell suspension was centrifuged, the supernatant removed, and the pellet resuspended in the according amount of cryomedia. Cryotubes were then frozen at - 80°C using a Mr. Frosty freezing container.

### 2.8 Determination of infarct size

The weight of the entire heart, the left ventricle after removal of the atria and the right ventricle, and each individual slice was recorded. In addition, the heart, the left ventricle, and each slice were photographed from both the apical and non-apical sides. This provided accurate, macroscopic documentation of the infarct and surrounding areas, as well as the healthy myocardium. Using the image processing software ImageJ, the area of infarcted tissue was manually measured in each slice—both apically and non-apically. The area of the entire slice, minus the area of the ventricle, was also measured. The infarct areas of the left ventricular slices were multiplied by the respective weight of each slice, summed, and used in conjunction with the weight of the entire heart to calculate the relative and absolute size of the infarct.

### 2.9 Hematoxylin-Eosin staining of porcine cardiac tissue

Thawed 10µm cryosections were fixed in 4% paraformaldehyde (PFA) for 30 minutes. The sections were subsequently stained with 0.1% Mayer’s hematoxylin (Morphisto, Offenbach, Germany) for 10 min, followed by bluing under running tap water for an additional 10 min. Counterstaining was performed using 0.3% eosin Y solution (Morphisto) for 5 min. After rinsing with distilled water (dH_2_O), the sections were dehydrated through a graded ethanol series, cleared in xylene followed by a xylene substitute (ROTI®Histol, Carl Roth, Karlsruhe, Germany), and mounted with a rapid mounting medium (Entellan, Merck, Darmstadt, Germany). The prepared slides were then examined using a light microscope (Keyence, BZ-X810, Frankfurt am Main, Germany).

### 2.10 Picrosirius Red (PSR) staining of porcine cardiac tissue

Thawed cryosections were fixed in 4% paraformaldehyde (PFA) for 30 min, then dipped 2–3 times in demineralized water followed by staining with 0.01% Picrosirius Red (PSR) solution (Morphisto, Offenbach, Germany) for 60 min. After rinsing with distilled water (dH_2_o), the sections were dehydrated through a graded ethanol series and cleared in xylene followed by a xylene substitute (ROTI® Histol, Carl Roth, Karlsruhe, Germany). Finally, the sections were mounted with a rapid mounting medium (Entellan, Merck, Darmstadt, Germany). Fluorescence microscopy was performed using a DMi8 microscope (Leica Microscopy, Wetzlar, Germany), enabling the differentiation of PSR-positive (collagen-rich) and PSR-negative tissue based on their distinct fluorescence properties.

### 2.11 Protein extraction and protein quantification

For protein extraction, myocardial tissue from the border zone and the infarct core, which had been frozen specifically for this purpose, was used. Approximately 20 mg of the frozen tissue was first washed twice in cold PBS to remove any blood. The myocardium was then placed in lysis buffer (radioimmunoprecipitation assay buffer (RIPA buffer) containing PMSF, phenylmethanesulfonyl fluoride powder, 0.5 mM, Sigma Aldrich, Merck KG, Darmstadt, Germany; and proteinase inhibitor, cOmplete™, EDTA-free Protease Inhibitor Cocktail Tablets, Millipore®, Midland Scientific Inc, La Vista, NE, USA) and briefly homogenized using a TissueRuptor (TissueRuptor Disposable Probes, Qiagen, Hilden, Germany). After one minute in the ultrasonic bath, the samples were incubated for 30 minutes with occasional mixing, and the clear supernatant was collected after centrifugation and frozen at -80°C. A small portion of the protein lysate was used directly for quantification via the bicinchoninic acid assay (BCA assay, Pierce™ BCA Protein Assay Kit, Thermo Fisher Scientific, Carlsbad, CA, USA). The BCA assay was performed according to the manufacturer’s instructions.

### 2.12 IL 1β – ELISA

Interleukine 1-beta was quantified by a Quantikine® ELISA porcine IL – 1β Immunoassay (R&D Systems Inc., Minneapolis, Canada) and the ELISA was carried out according to the manufacturer’s instructions.

### 2.13 RNA isolation and sequencing

Frozen myocardial samples were used for RNA isolation. RNA isolation was performed using the RNeasy Mini Kit (RNeasy Kits for RNA Purification, Qiagen, Hilden, Germany) according to the manufacturer’s instructions. In addition, Safe-Lock Biopur® tubes (1.5 ml & 2 ml, Eppendorf, Germany) were used. RNA quality was checked using a 5200 Fragment Analyzer with the DNF-471-33 - SS Total RNA 15nt kit (Agilent Technologies). The RQN for all samples was ≥ 7.2. DNA libraries suitable for sequencing were prepared from 300 ng of total RNA with oligo-dT capture beads for poly-A-mRNA enrichment using the Watchmaker mRNA Library Preparation Kit (Watchmaker Genomics) according to manufacturer’s instructions (½ volume). After 11 cycles of PCR amplification, the size distribution of the barcoded DNA libraries was estimated ∼335 bp by electrophoresis on 5200 Fragment Analyzer with the DNF-474-33 - HS NGS Fragment 1-6000bp kit (Agilent Technologies). Sequencing of pooled libraries, spiked with 1% PhiX control library, was performed at 33-38 million reads/sample in single-end mode with 100 nt read length on the NextSeq 2000 platform (Illumina). Demultiplexed FASTQ files were generated with bcl-convert v4.3.6 (Illumina).

To assure high sequence quality, Illumina reads were quality- and adapter-trimmed via Cutadapt (11) v2.5 using a cutoff Phred score of 20 in NextSeq mode, and reads without any remaining bases were discarded (parameters: --nextseq-trim=20 -m 1 -a AGATCGGAAGAGCACACGTCTGAACTCCAGTCAC). Processed reads were subsequently mapped to the pig genome (NCBI RefSeq assembly GCF_000003025.6/Sscrofa11.1) using STAR (12) v2.7.2b with default parameters but including transcript annotations from RefSeq annotation version 106 for Sscrofa11.1. This annotation was also used to generate read counts on exon level summarized for each gene via featureCounts v1.6.4 from the Subread package (13). Multi-mapping and multi-overlapping reads were counted strand-specific and reversely stranded with a fractional count for each alignment and overlapping feature (parameters: -s 2 -t exon -M -O --fraction). The count output was utilized to identify differentially expressed genes using DESeq2 (14) v1.24.0. Read counts were normalized by DESeq2 and fold-change shrinkage was conducted by setting the parameter betaPrior to TRUE. Differential expression was assumed at adjusted p-value after Benjamini-Hochberg correction (padj) <0.05 and |log2FoldChange| ≥ 1. clusterProfiler (15) v4.8.1 was used to perform functional enrichment analyses based on Gene Ontology (GO) terms and the “Reactome inflammasomes” pathway (R-HSA-622312), which was mapped to the pig annotation via gene name matching. The GSEA function was applied for gene set enrichment analysis considering the DESeq2 log2FoldChange of all analyzed genes.

### 2.14 Statistical data analysis

Statistical analysis and data visualization were performed using GraphPad Prism v10.6.0 (GraphPad Software, San Diego, CA, USA). Values for each animal are shown individually with horizontal bars indicating means per group. The individual statistical tests applied are indicated in the figure legends. A p-value < 0.05 was considered statistically significant.

## 3. Results

Nineteen landrace pigs underwent occlusion of the left anterior descending coronary artery for 90 minutes to induce myocardial infarction (MI). 13 animals survived the procedure and were randomly assigned to oral treatment with 0.05 mg/kg colchicine or placebo on day 3 post MI. All animals were treated per protocol and survived until analysis, i.e. day 7 after MI (Fig. 1A). The relative infarct size determined as infarct area/ left ventricular weight also did not significantly differ between treatment groups (Fig.1B, C). Neither echocardiography nor necropsy revealed left ventricular (LV) thrombi. Histology showed evolved replacement of dead myocardial tissue by a de novo formed collagen matrix and intensive infiltration of leukocytes without obvious differences between treatment groups at this point of analysis (Fig.1D-G).

**Figure 1:**
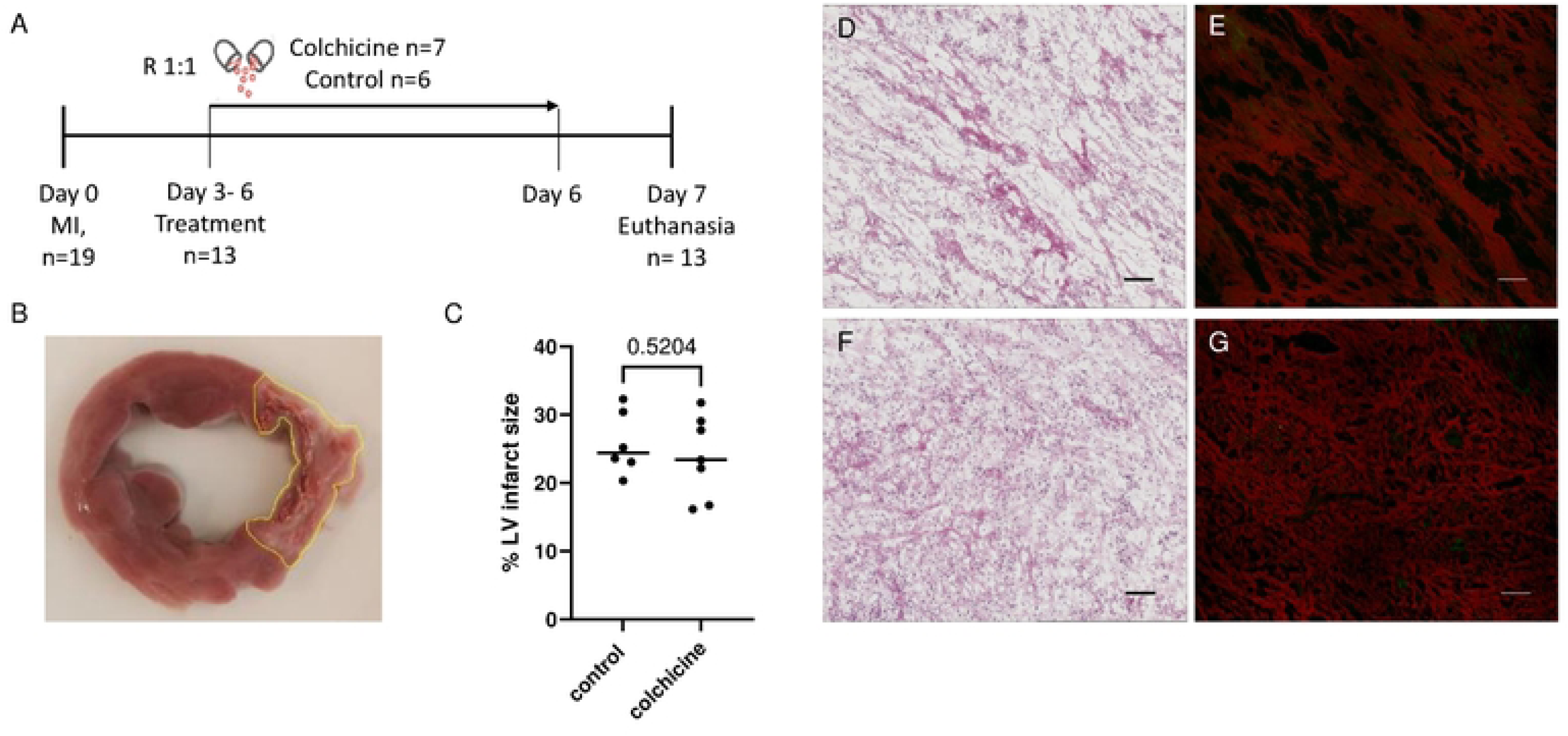
Ischemia-reperfusion model (MI). A: Study scheme B: Representative picture of a left ventricular slice including the pale infarct area marked by a yellow line at day 7. C: Infarct weight relative to LV weight (g/g) at day 7. D-G: Histological sections of the infarct zone of colchicine (D, E) and control animals (F, G) stained by H.E. (D, F), and PSR (E, G), bar represents 100µm. p values were determined by an unpaired t-test.

Echocardiography was performed before MI and on day 7 (Fig. 2A). The EF was significantly lower in the colchicine group, but the stroke volumes, end-diastolic diameters and E/E‘, reflecting diastolic function, on day 7 were statistically not different. The end-diastolic diameter at day 7 and its change from baseline, reflecting early left-ventricular (LV) remodelling, were also not different between treatment groups (Fig. 2B). However, we observed no significant difference regarding the change (day 7-baseline before MI) of ejection fraction (EF) and end-diastolic diameter between treatment groups (Fig. 2C).

**Figure 2:**
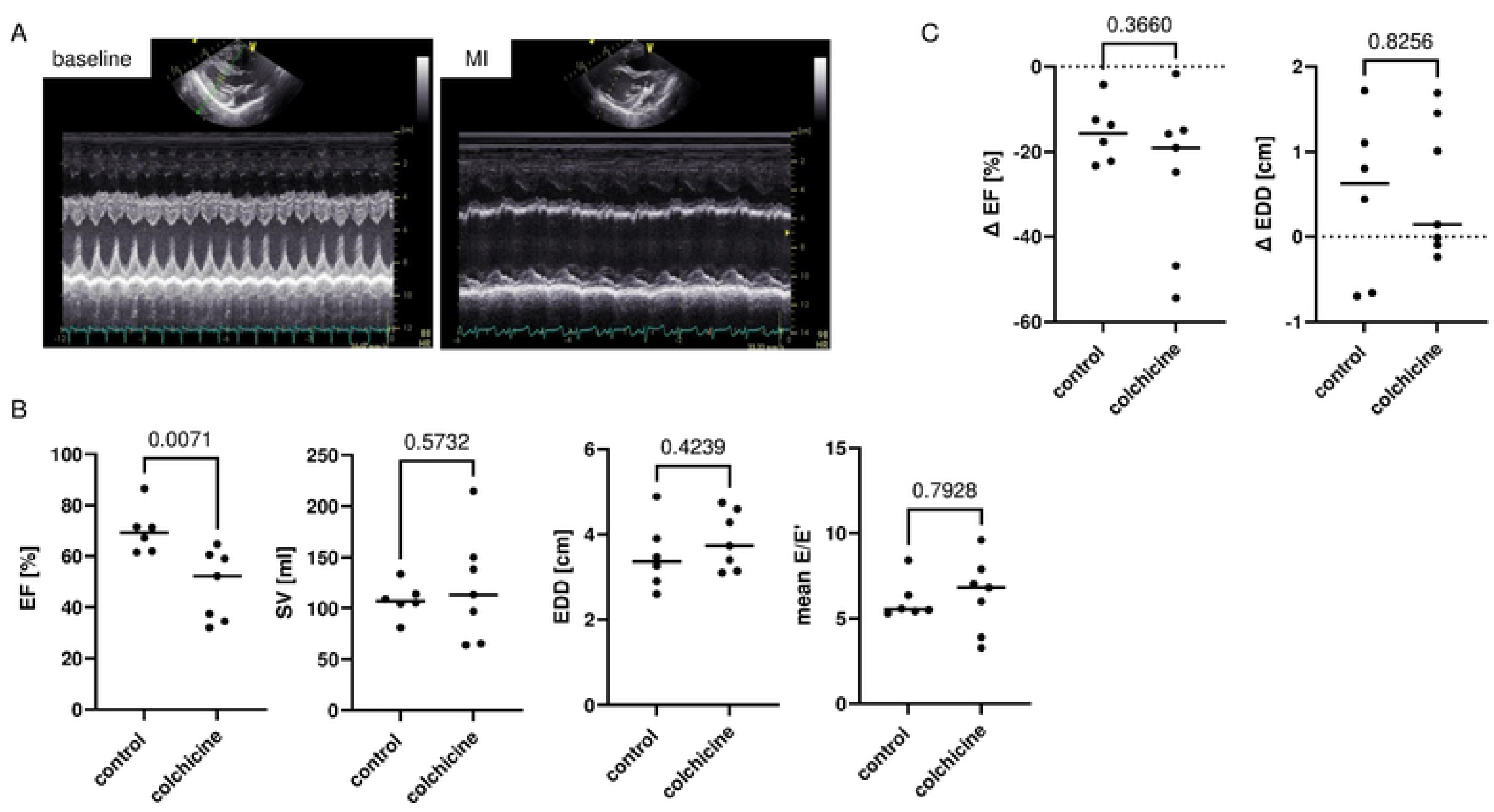
Echocardiographic analysis A: M-mode of long axis-views show representative longitudinal analysis of the left ventricle before and after MI (day 7). B: EF, stroke volume (SV), EDD, and E/E‘ reflecting diastolic function on day 7. p values were determined by Mann-Whitney tests. C: change in ejection fraction (EF) and end-diastolic diameter (EDD) between baseline and day 7.

As rodent studies indicated that colchicine modifies leukocyte recruitment and differentiation in the injured myocardium, we analysed the leukocyte composition of the infarct, infarct border and remote zone by flow cytometry. Our panel allowed for identification of myeloid cells, including macrophages and neutrophils (Fig. 3A-C, gating strategy in Suppl. Fig. I). NLRP3 expression in, both, macrophages (Fig. 3D) and neutrophils (Fig. 3E) was similar in both groups (Fig. 3D, E). High CD163 expression allows to identify anti-inflammatory and tissue-resident macrophages. Macrophage subsets defined by CD163 surface expression level were not significantly different between treatment groups. The distribution of lymphocyte and T cell subsets, including conventional CD4^+^, regulatory Foxp3^+^ and cytotoxic CD8b^+^ T-cells, also were not different between the treatment groups (Fig. 4A-E, gating strategy in Suppl. Fig. II).

**Figure 3:**
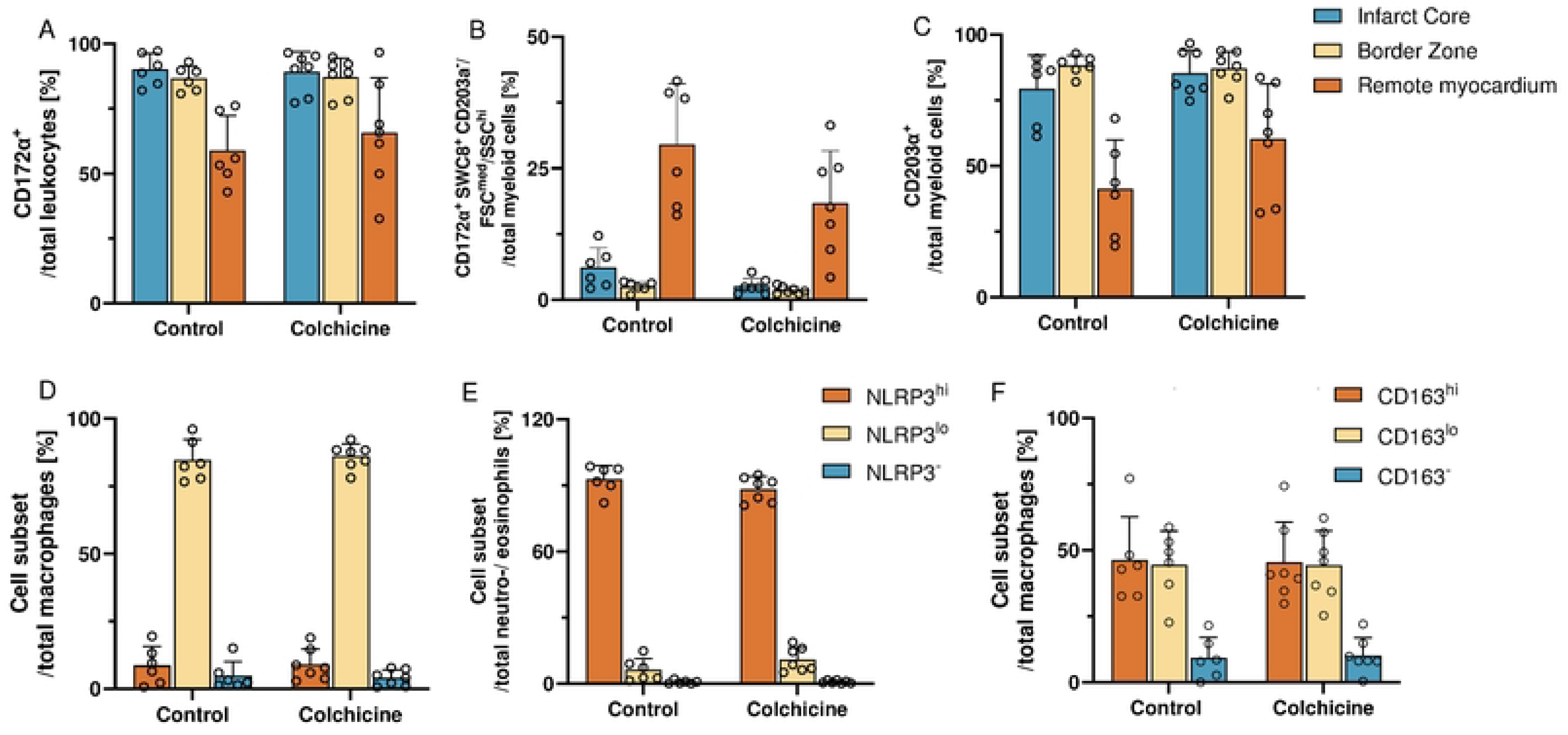
Flow-cytometric analysis of myeloid cells within the myocardium. A: Myeloid cells among total leukocytes B: Neutrophils C: Macrophages D: NLRP3 expression on macrophages within infarct zone E: NLRP3 expression on neutrophils and eosinophils within infarct zone F: CD163 expression on macrophages within the infarct zone.

**Figure 4:**
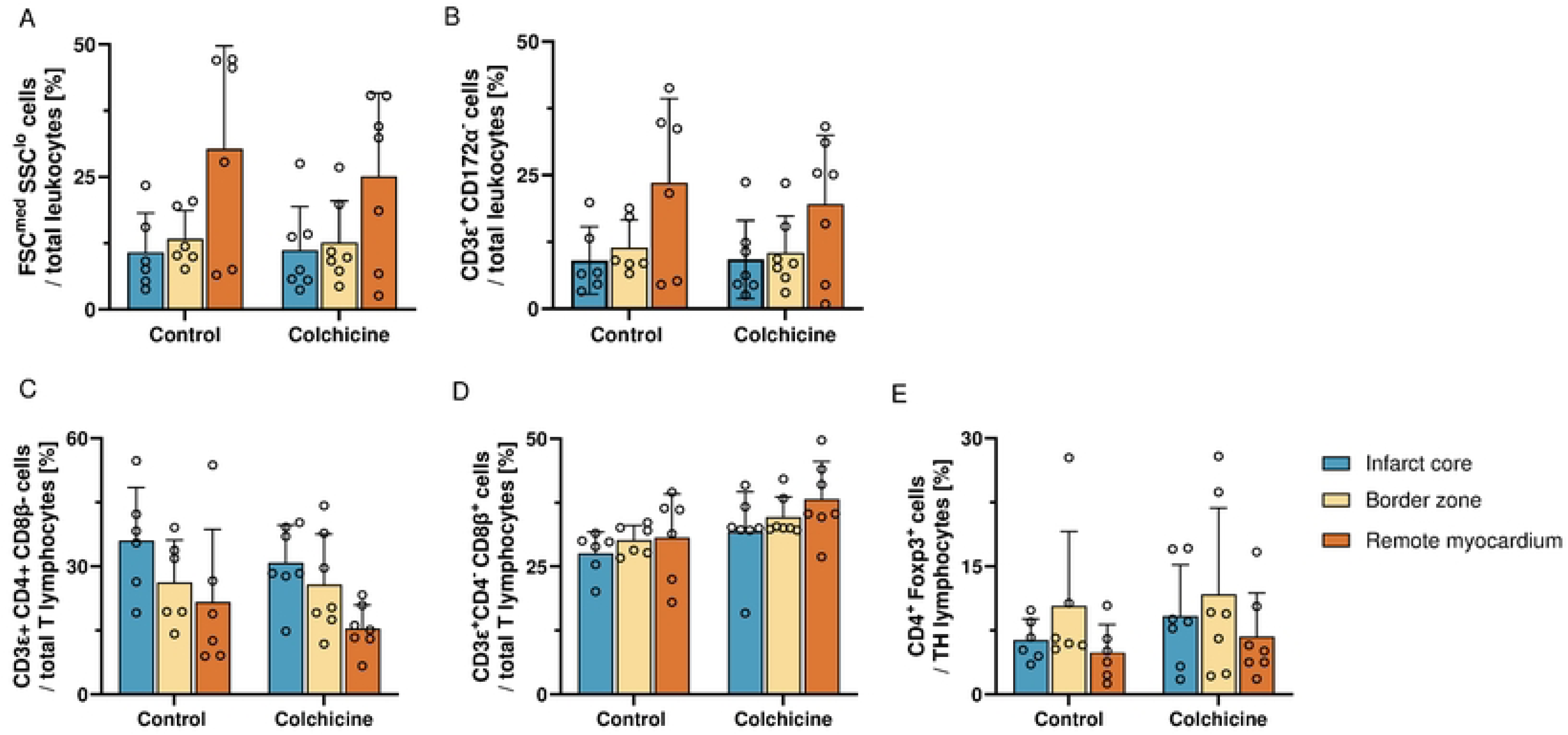
Flow-cytometric analysis of lymphoid cells within the myocardium. A: Lymphoid cells among all leukocytes. B: T-cells C: CD4^+^ T-cells. D: CD8^+^ T-cells E: Foxp3^+^ CD4^+^ T–cells.

Because a major anti-inflammatory mechanism of colchicine is inhibition of the NLRP3 inflammasome, which cleaves pro-IL-1β prior to secretion, we quantified IL-1β levels in the infarct and infarct border zone. The analysis revealed no significant differences between treatment groups (Fig. 5A, B).

**Figure 5:**
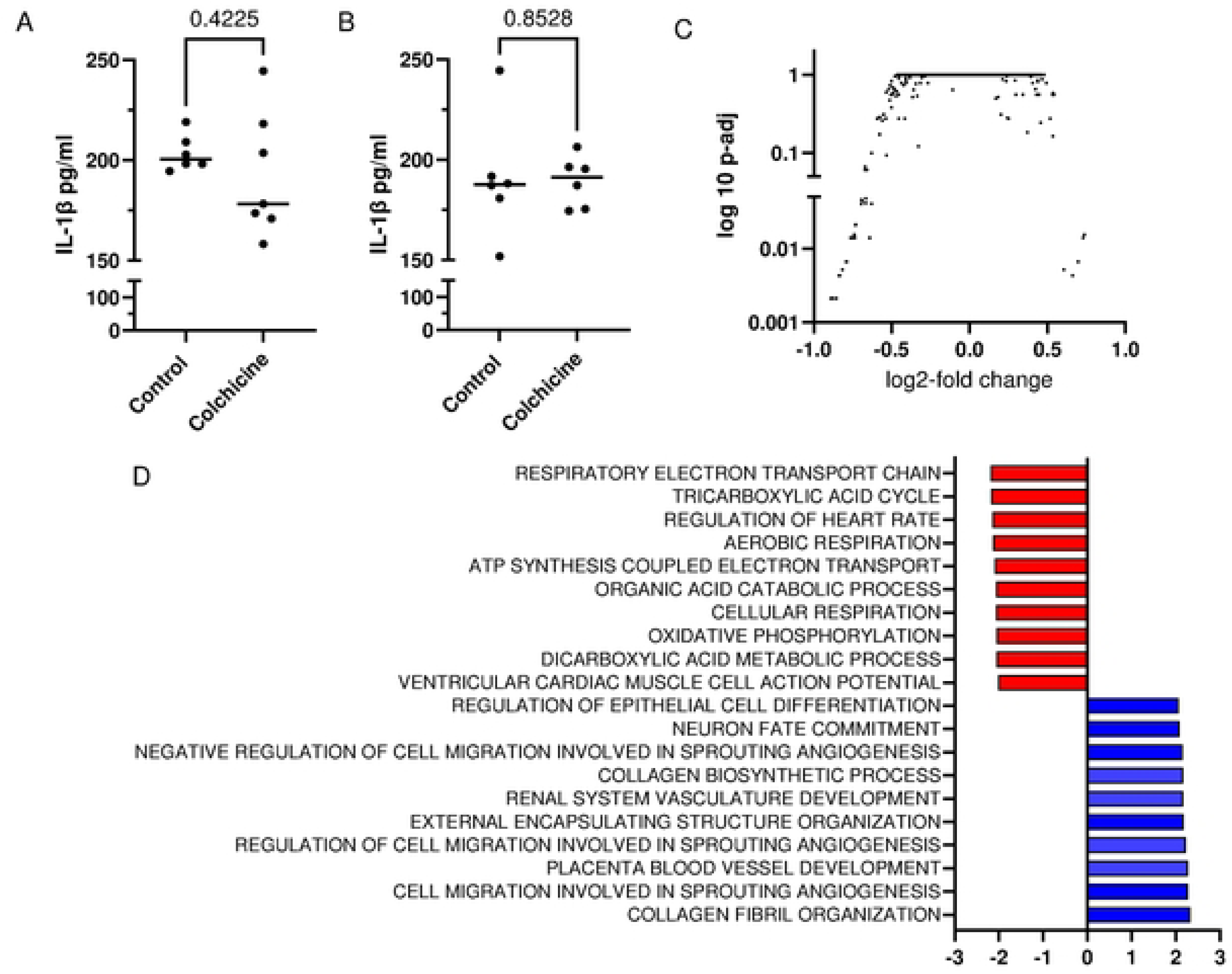
Gene expression analysis A: Interleukin-1β within the infarct zone and B: within the border zone. p values were determined by Mann-Whitney tests. C: Bulk-RNA-sequencing colchicine vs. control D: NES for top regulated GO terms colchicine vs. control, p-adjusted <0.05 each. Red marks downregulated and blue upregulated gene sets in colchicine group.

To get a broad and unbiased picture on how colchicine treatment might have changed the transcriptional landscape in the infarct zone, we employed bulk-RNA-sequencing. Within the infarct zone, few single transcripts were differentially expressed (Fig. 5C, Suppl. File I). Unbiased exploratory gene set enrichment analysis (GSEA) revealed GO terms representing collagen metabolism and angiogenesis to be upregulated in colchicine treated animals whereas GO terms reflecting energy metabolism, oxidative phosphorylation and cardiomyocyte function were downregulated (Fig.5D, Suppl. File II). Notably, gene sets representing inflammation were not significantly enriched in the control versus the Colchicine group. To test more specifically if the transcriptome revealed a treatment dependent inflammasome-signalling signature, we performed GSEA using the gene set “reactome inflammasomes”. It revealed no significant enrichment in the control versus the Colchicine group (NES: -0.97, p-value = 0.49). Individual transcripts of this gene set also showed no differential regulation (Suppl. Fig III). Together, colchicine treatment induced no significant impact on proinflammatory transcriptional activity.

## 4. Discussion

Our randomised placebo-controlled preclinical study in pigs demonstrated that oral colchicine treatment produced, overall, no modulation of leukocyte composition and transcription of inflammatory genes in the reperfused myocardium. Moreover, treatment neither reduced infarct size nor preserved systolic function. Notably, treatment was well tolerated and no safety concerns were identified. In particular, no left ventricular thrombi were observed.

These observations need to be interpreted in the context of a recent clinical trial (8) evaluating oral colchicine treatment in patients with acute MI. When colchicine was administered before or right after initiation of reperfusion, more left ventricular thrombi were observed but there was no beneficial effect on cardiac injury or preservation of systolic function. Hence, delayed initiation of treatment may avoid the generation of thrombi but is likewise ineffective regarding preservation of systolic function.

How can the discrepancies in efficacy between several promising small animal studies (4–7) and the large animal and clinical studies be explained? The proposed cardioprotective mechanisms of colchicine in reperfused MI include attenuation of myeloid cell infiltration and suppression of NLRP3 inflammasome-mediated interleukin-1β release with subsequent pro-inflammatory signalling affecting many cell types. As neither leukocyte composition nor IL-1β protein levels or global patterns of proinflammatory transcription differed between treatment groups, the most likely explanation for the lack of efficacy is that the treatment regimen failed to adequately suppress cardiac inflammation on day 7 post MI. The colchicine concentrations achieved within the myocardium were likely insufficient to effectively inhibit NLRP3 activity and leukocyte infiltration, effects that were reported to attenuate cardiac injury in rodent models (4–7). Notably, administration of a tenfold higher dose in a dose-finding experiment resulted in clinically relevant gastrointestinal side effects, which led us to select the current dose. It equals the dose previously studied in pigs (16) and is approximately tenfold higher than the doses approved for secondary prevention in patients after MI. However, owing to interspecies differences in metabolism, it cannot be assumed that doses in humans and pigs are biologically equivalent. Higher doses of colchicine have been used in rodents, and both the pharmacokinetic and pharmacodynamic effects may differ owing to species-specific differences in metabolism. Moreover, in rodent models, colchicine was administered mostly before or immediately at the onset of ischaemia or reperfusion, targeting the earliest phases of reperfusion. Our gene expression data revealed that colchicine modulated transcriptional activity related to angiogenesis and collagen metabolism. This finding corresponds to the observation of reduced collagen deposition in the aforementioned pig study evaluating colchicine treatment (16). GO terms reflecting energy metabolism, oxidative phosphorylation and cardiomyocyte function were downregulated in the colchicine group. This might have contributed to the significantly lower EF in the colchicine group albeit equal infarct sizes. However, considering that the change in EF was not significantly different, this was probably more driven by differences in baseline EF.

As inflammation after MI is not purely deleterious; a carefully orchestrated sequence of inflammatory and reparative responses is essential for clearance of necrotic tissue and formation of a stable scar. If colchicine is given too early or at a dose that excessively suppresses key inflammatory pathways, it might even impair effective healing, alter scar composition, and potentially favour adverse remodelling. Notably, the increased prevalence of left ventricular thrombi in patients treated with colchicine in acute MI is reminiscent of the effect observed following macrophage depletion in experimental MI (17). We selected day 7 for analysis because, based on our previous findings, this time point corresponds to the transition from the acute inflammatory phase towards the activation of anti-inflammatory and reparative programmes in leukocytes (18). However, neither flow cytometric profiling nor transcriptional analyses provided evidence of suppression of pro-inflammatory processes at this stage. It remains possible, however, that single-cell RNA sequencing could reveal more subtle colchicine-induced changes in leukocyte differentiation and functional states that are not captured by the bulk transcriptional or flow cytometric approaches applied.

We acknowledge the limitation that echocardiography for the assessment of left ventricular function and geometry in pigs has its limitations. Furthermore, both the treatment duration and follow-up period were relatively short. Consequently, we cannot exclude the possibility that analyses performed after a longer treatment period or at a later follow-up time point might have detected effects on inflammation that were not apparent at the selected time of assessment. Nevertheless, our findings are consistent with those of another porcine study, in which the same daily dose of colchicine likewise failed to produce significant effects on infarct size or systolic function, as assessed by MRI at 30 days following MI (16). Hence, it is rather unlikely that a longer period of treatment might have induced significant protection.

The biology of chronic plaque inflammation in coronary artery disease differs substantially from that of acute ischaemic injury and the subsequent healing response within the myocardium. The totality of evidence from large clinical trials testing colchicine for secondary prevention, including meta-analyses indicate significant effects of colchicine on vascular events and ischaemia-driven revascularisation, albeit without a reduction in all-cause or cardiovascular mortality (19, 20). Mechanistically, at the dose used for secondary prevention, colchicine is effective in attenuating chronic, low-grade vascular inflammation and in preventing plaque destabilisation. However, at the doses tested in pig and human studies, its suppressive effect on the acute, high-intensity inflammatory response within the infarcted myocardium is likely insufficient. Given its toxicity profile, studies employing substantially higher doses, which might achieve more effective suppression of myocardial inflammation after MI, are not feasible. As an alternative approach, specific inflammasome inhibitors currently seem to be more promising therapeutic agents, both for preserving cardiac function after MI and for preventing subsequent ischaemic events (21). Of note, treatment with the compound MCC950 lowered myocardial IL-1β levels, reduced infarct size and preserved systolic function in a pig model of myocardial infarction (22).

Collectively, colchicine has an established and guideline-supported role in secondary prevention of ischaemic events for patients after MI. Preclinical rodent studies suggested additional potential benefits in mitigating ischemia–reperfusion injury and preserving systolic function after MI, but these have not been corroborated by a clinical trial and large-animal studies. This may reflect insufficient suppression of inflammatory response within the infarcted myocardium at clinically tolerable colchicine doses.

## 6. Acknowledgments

We greatly appreciate the technical support of Katja Blouin, Nicole Maier, Sandra Werner-Wittig and Jutta Meißner-Weigl for their outstanding and supportive work in the laboratory as well as Verena Burkard and Kerstin Körner for their invaluable help with animal experiments especially for their dedicated animal care. We thank the Core Unit Systems Medicine at the University of Würzburg for excellent technical support and RNA-seq data generation and Prof. Uwe Müller, for the valuable supervision of Rebekka Grampp’s thesis.

---

**Supplemental Figure I:**
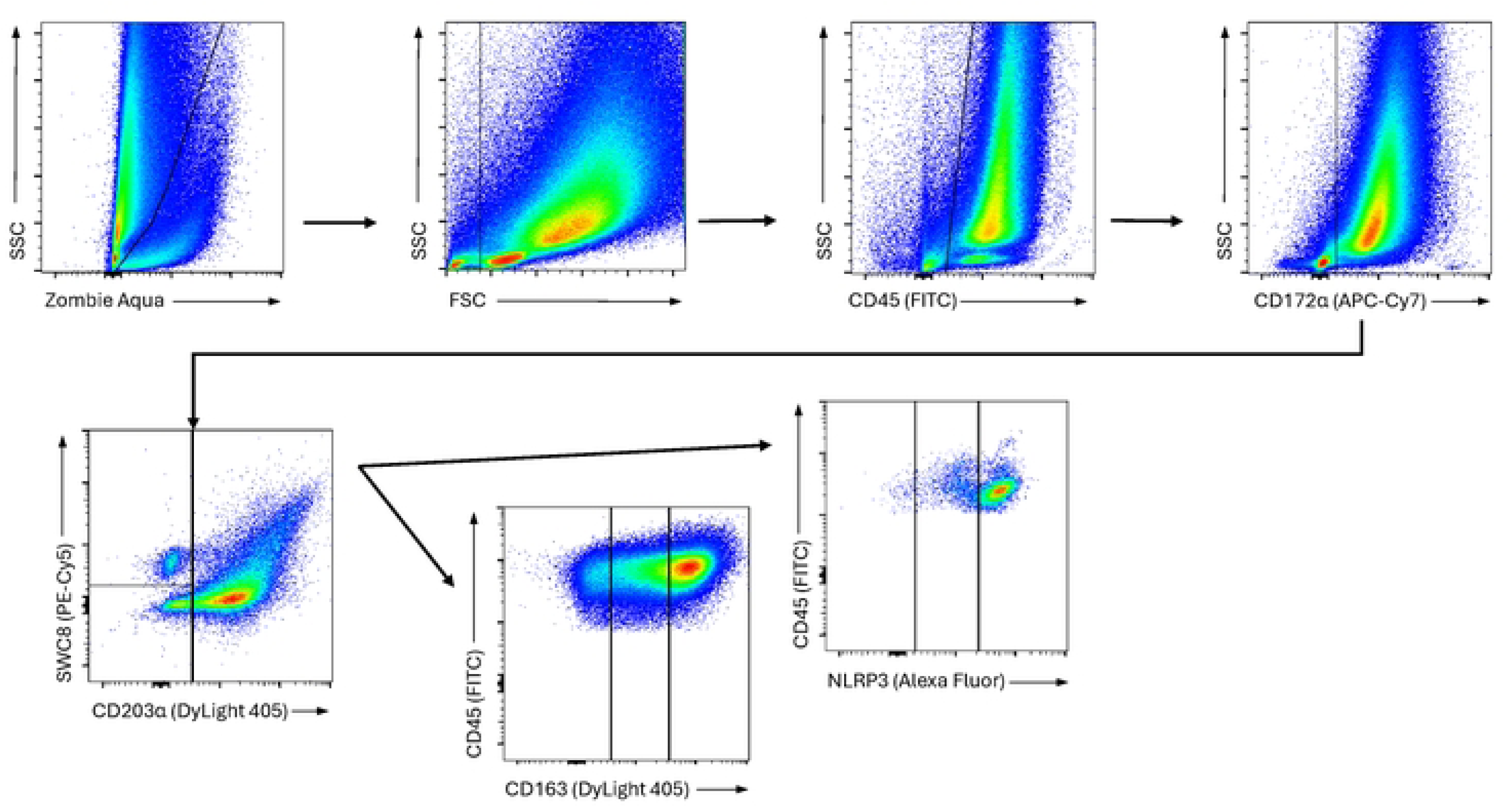
Gating strategy of the myeloid panel

**Supplemental Figure II:**
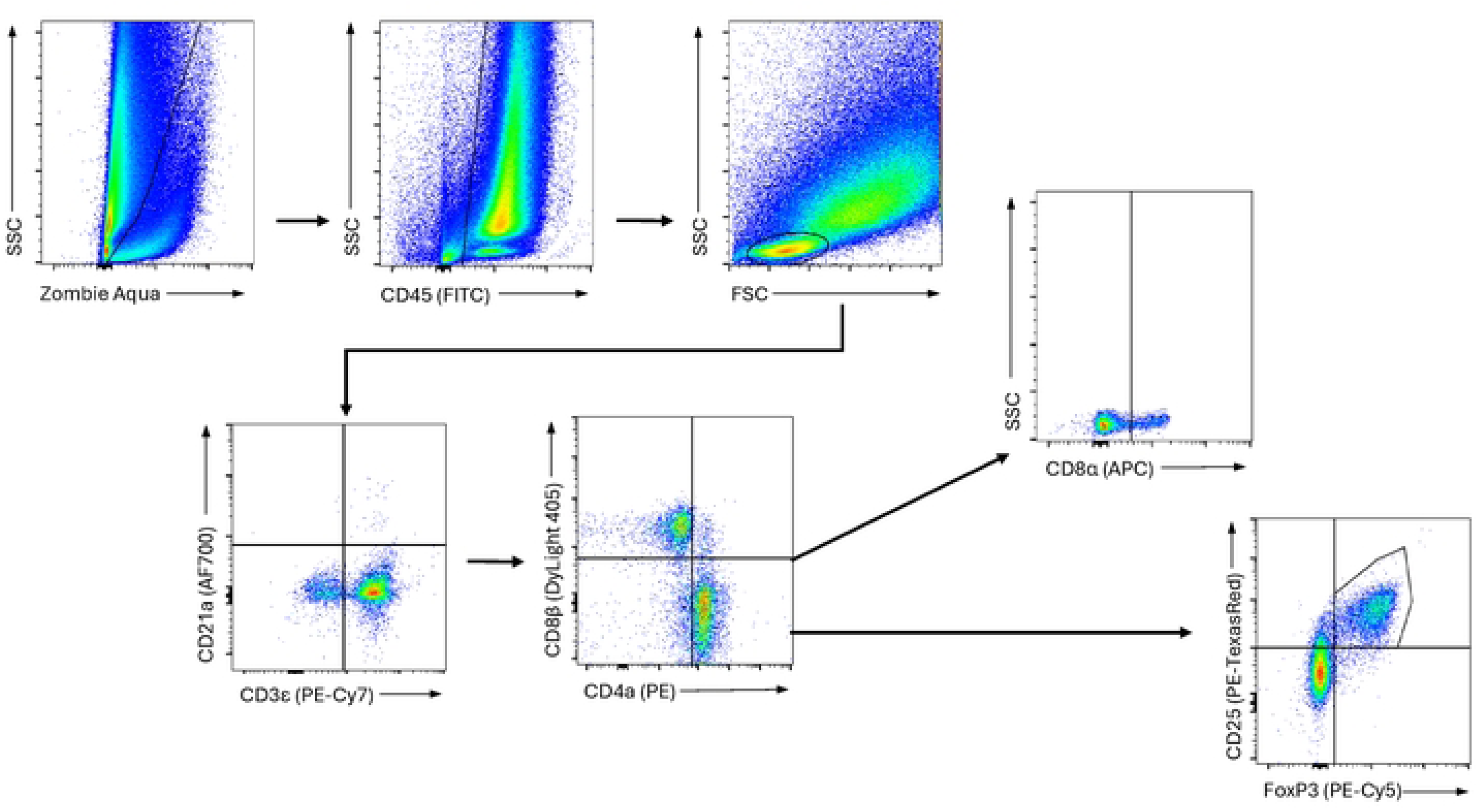
Gating strategy of the lymphoid panel

**Supplemental Figure III:**
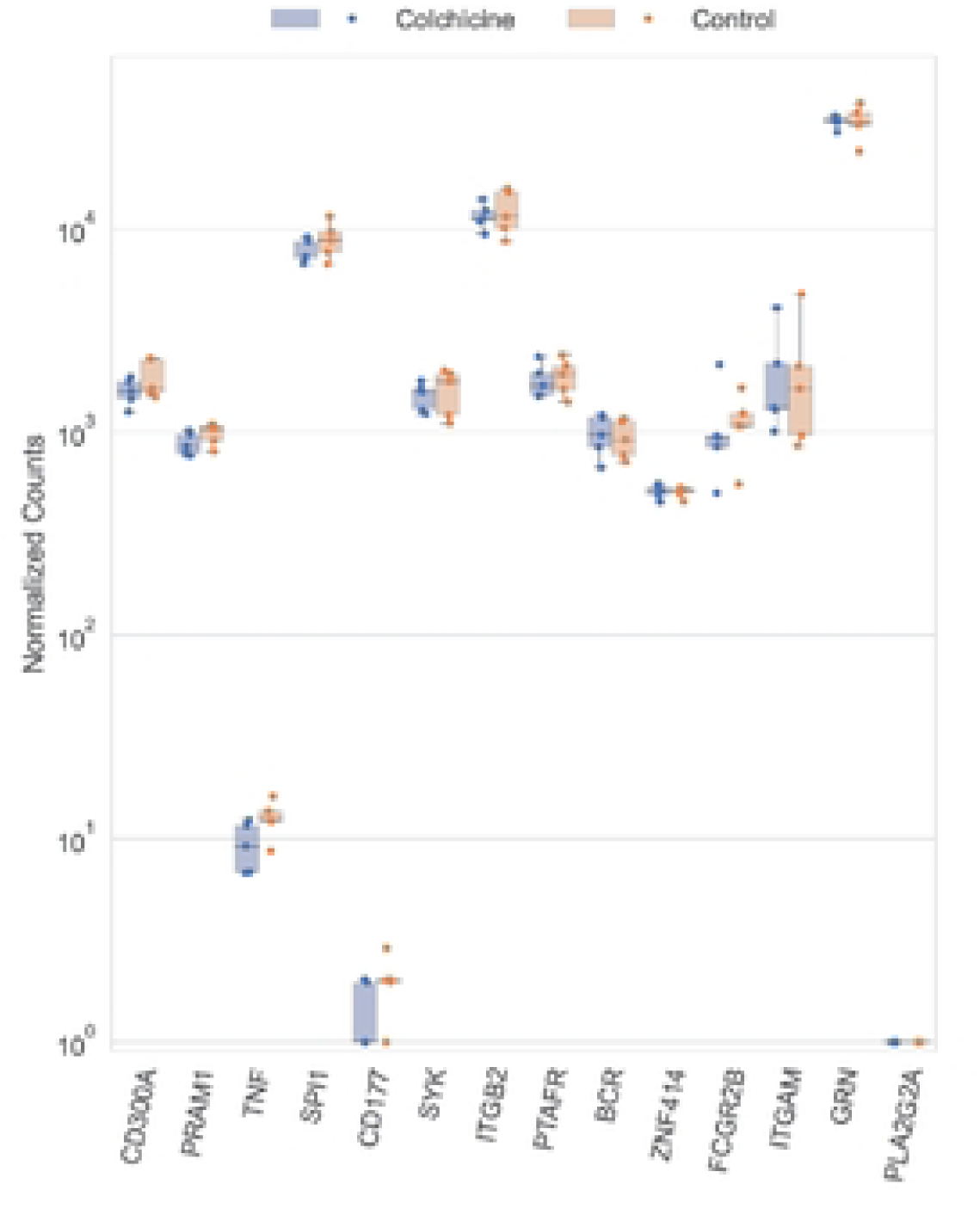
Individual gene expression within gene set „Reactome inflammasomes“(R-HAS-622312). Condition-specific expression levels for the Reactome Inflammasomes gene set were plotted using the Python package seaborn* v0.13.2. An overlay of boxplots and stripplots was applied to depict data points and distribution for DESeq-normalized counts on a log-scale y-axis for each gene on the x-axis.

**Supplemental table I:** List of Monoclonal primary antibodies used for flow cytometry (FC) analysis of heart samples.

|  | Antigen | Conjugate | Isotype | Clone | Reactivity | Format | Source | Concentration |
|---|---|---|---|---|---|---|---|---|
| All cells | CD45 | FITC | IgG1 | K252.1E4 | mouse:pig | labelled | Biorad | 1:50 |
| Myeloid cells | CD203a | DyLight 405 | IgG1 | PM18-7 | mouse:pig | unlabelled | Biorad | 1:200 |
|  | CD163 | Pe-Texas Red | IgG1 | 2A10/11 | mouse:pig | unlabelled | Biorad | 1:200 |
|  | CD172α | APC-Cy7 | IgG2b | 74-22-15a | mouse:pig | unlabelled | BD | 1:200 |
|  | SWC8 | / | IgM | MIL3 | mouse:pig | unlabelled | Thermo Fisher | 1:20 |
|  | GPNMB | PE | IgG | polyklonal | rabbit:pig | labelled | Bioss | 1:100 |
|  | NLRP3 | AF 647 | IgG | polyklonal | rabbit:pig | labelled | Bioss | 1:200 |
|  | mIgM | PE-Cy5 | IgG2a | II/41 | rat:mouse | labelled | Thermo Fisher | 1:200 |
|  | FoxP3 | PE-Cy5 | IgG2a | FJK-16s | rat:mouse | labelled | Thermo Fisher | 1:50 |
| Lymphoid cells | CD25 | PE-Texas Red | IgG1 | K231.3B2 | mouse:pig | unlabelled | Biorad | 1:100 |
|  | CD172α | APC-Cy7 | IgG2b | 74-22-15a | mouse:pig | unlabelled | BD | 1:200 |
|  | CD8α | APC | IgG2a | 76-2-11 | mouse:pig | labelled | Thermo Fisher | 1:200 |
|  | CD8β | DyLight 405 | IgG1 | PPT23 | mouse:pig | unlabelled | Biorad | 1:200 |
|  | CD3ε | PE-Cy7 | IgG2a | BB23-8E6-8C8 | mouse:pig | labelled | BD | 1:200 |
|  | CD21 | AF 700 | IgG1 | BB6-11C9.6 | mouse:pig | labelled | Novus Biologicals | 1:200 |
|  | CD4α | PE | IgG2b | 74-12-4 | mouse:pig | labelled | BD | 1:200 |

## References

1. Leung YY, Yao Hui LL, Kraus VB. Colchicine--Update on mechanisms of action and therapeutic uses. Semin Arthritis Rheum. 2015;45(3):341–50.

2. Nidorf SM, Fiolet ATL, Mosterd A, Eikelboom JW, Schut A, Opstal TSJ, et al. Colchicine in Patients with Chronic Coronary Disease. N Engl J Med. 2020;383(19):1838–47.

3. Tardif JC, Kouz S, Waters DD, Bertrand OF, Diaz R, Maggioni AP, et al. Efficacy and Safety of Low-Dose Colchicine after Myocardial Infarction. N Engl J Med. 2019;381(26):2497– 505.

4. Fujisue K, Sugamura K, Kurokawa H, Matsubara J, Ishii M, Izumiya Y, et al. Colchicine Improves Survival, Left Ventricular Remodeling, and Chronic Cardiac Function After Acute Myocardial Infarction. Circ J. 2017;81(8):1174–82.

5. Bakhta O, Blanchard S, Guihot AL, Tamareille S, Mirebeau-Prunier D, Jeannin P, et al. Cardioprotective Role of Colchicine Against Inflammatory Injury in a Rat Model of Acute Myocardial Infarction. J Cardiovasc Pharmacol Ther. 2018;23(5):446–55.

6. Wang L, Peng Y, Song L, Xia D, Li C, Li Z, et al. Colchicine-Containing Nanoparticles Attenuates Acute Myocardial Infarction Injury by Inhibiting Inflammation. Cardiovasc Drugs Ther. 2022;36(6):1075–89.

7. Mori H, Taki J, Wakabayashi H, Hiromasa T, Inaki A, Ogawa K, et al. Colchicine treatment early after infarction attenuates myocardial inflammatory response demonstrated by (14)C-methionine imaging and subsequent ventricular remodeling by quantitative gated SPECT. Ann Nucl Med. 2021;35(2):253–9.

8. Mewton N, Roubille F, Bresson D, Prieur C, Bouleti C, Bochaton T, et al. Effect of Colchicine on Myocardial Injury in Acute Myocardial Infarction. Circulation. 2021;144(11):859– 69.

9. 9. Jolly SS, d’Entremont MA, Lee SF, Mian R, Tyrwhitt J, Kedev S, et al. Colchicine in Acute Myocardial Infarction. N Engl J Med. 2025;392(7):633–42.

10. Bouabdallaoui N, Tardif JC, Waters DD, Pinto FJ, Maggioni AP, Diaz R, et al. Time-to-treatment initiation of colchicine and cardiovascular outcomes after myocardial infarction in the Colchicine Cardiovascular Outcomes Trial (COLCOT). Eur Heart J. 2020.

11. Martin M. Cutadapt removes adapter sequences from high-throughput sequencing reads. EMBnetjournal. 2011.

12. Dobin A, Davis CA, Schlesinger F, Drenkow J, Zaleski C, Jha S, et al. STAR: ultrafast universal RNA-seq aligner. Bioinformatics. 2013;29(1):15–21.

13. Liao Y, Smyth GK, Shi W. featureCounts: an efficient general purpose program for assigning sequence reads to genomic features. Bioinformatics. 2014;30(7):923–30.

14. Love MI, Huber W, Anders S. Moderated estimation of fold change and dispersion for RNA-seq data with DESeq2. Genome Biol. 2014;15(12):550.

15. Wu T, Hu E, Xu S, Chen M, Guo P, Dai Z, et al. clusterProfiler 4.0: A universal enrichment tool for interpreting omics data. Innovation (Camb). 2021;2(3):100141.

16. Aimo A, Martinez-Falguera D, Barison A, Musetti V, Masotti S, Morfino P, et al. Colchicine added to standard therapy further reduces fibrosis in pigs with myocardial infarction. J Cardiovasc Med (Hagerstown). 2023;24(11):840–6.

17. Frantz S, Hofmann U, Fraccarollo D, Schafer A, Kranepuhl S, Hagedorn I, et al. Monocytes/macrophages prevent healing defects and left ventricular thrombus formation after myocardial infarction. FASEB J. 2013;27(3):871–81.

18. Schnitter F, Stangl F, Noeske E, Bille M, Stadtmuller A, Vogt N, et al. Characterizing the immune response to myocardial infarction in pigs. Basic Res Cardiol. 2024;119(3):453–79.

19. d’Entremont MA, Poorthuis MHF, Fiolet ATL, Amarenco P, Boczar KE, Buysschaert I, et al. Colchicine for secondary prevention of vascular events: a meta-analysis of trials. Eur Heart J. 2025;46(26):2564–75.

20. Ebrahimi F, Ebrahimi R, Beer M, Schonenberger CM, Ewald H, Briel M, et al. Colchicine for the secondary prevention of cardiovascular events. Cochrane Database Syst Rev. 2025;11(11):CD014808.

21. Mo B, Ding Y, Ji Q. NLRP3 inflammasome in cardiovascular diseases: an update. Front Immunol. 2025;16:1550226.

22. van Hout GP, Bosch L, Ellenbroek GH, de Haan JJ, van Solinge WW, Cooper MA, et al. The selective NLRP3-inflammasome inhibitor MCC950 reduces infarct size and preserves cardiac function in a pig model of myocardial infarction. Eur Heart J. 2017;38(11):828–36.

